# Dosage compensation in *Bombyx mori* is achieved by partial repression of both Z chromosomes in males

**DOI:** 10.1101/2021.07.18.452821

**Authors:** Leah F. Rosin, Yang Chen, Elissa P. Lei

## Abstract

Interphase chromatin is organized precisely to facilitate accurate gene expression. The structure-function relationship of chromatin is epitomized in sex chromosome dosage compensation (DC), where sex-linked gene expression is balanced between males and females via sex-specific alterations to 3D chromosome structure. Studies in ZW-bearing species suggest that DC is absent or incomplete in most lineages except butterflies and moths, where male (ZZ) chZ expression is reduced by half to equal females (ZW). However, whether one chZ is inactivated (as in mammals) or both are partially repressed (as in *C. elegans*) is unknown. Using Oligopaints in the silkworm, *Bombyx mori*, we visualize autosome and chZ organization in somatic cells from both sexes for the first time. We find that *B. mori* interphase chromosomes are highly compact relative to *Drosophila* chromosomes. Importantly, we show that in *B. mori* males, both chZs are similar in size and shape and are more compact than autosomes or the female chZ after DC establishment, suggesting that both male chZs are partially and equally downregulated. We also find that in the early stages of DC, the female chZ repositions toward the nuclear center concomitant with increased Z-linked gene expression, revealing the first non-sequencing-based support for Ohno’s hypothesis. These studies represent the first visualization of interphase genome organization and chZ structure in Lepidoptera. We uncover striking similarities between DC in *B. mori* and *C. elegans*, despite these lineages harboring evolutionarily distinct sex chromosomes (ZW/XY), suggesting convergent evolution of DC mechanisms and a possible role for holocentricity in DC evolution.

**Significance:** Genes on sex chromosomes (for example, the X chromosome in humans) are regulated by distinct processes that do not affect non-sex chromosomes (autosomes). The expression of genes of sex chromosomes needs to be balanced between males and females due to differences in sex chromosome dosage (XY versus XX). This sex-specific gene regulation is called dosage compensation (DC). In most species, DC is achieved by altering the shape and compaction of sex chromosomes specifically in one sex. In this study, we use a chromosome painting approach called Oligopaints to examine how DC is achieved in silkworms for the first time. This is the first study to visualize this phenomenon in a species with ZW sex chromosomes, which evolved completely independently of XY. We find evidence supporting a long-standing model for how DC mechanisms evolved across all organisms, and we show high similarity between DC in silkworms and nematodes, suggesting these mechanisms have emerged multiple independent times throughout evolution.

## Introduction

The interphase genome is a complex network of 3D chromatin interactions. Individual chromosomes occupy discrete regions of the nucleus termed chromosome territories (CTs) (1) that are non-randomly organized in interphase. Chromosome size can influence CT position, with smaller chromosomes being more central in human nuclei (2), and studies from flies to humans have shown that gene-poor CTs are positioned closer to the nuclear periphery than gene-rich CTs (3–5). In agreement, both genomics and imaging-based approaches have revealed that the nuclear periphery is dense with repressive heterochromatin, while the nuclear interior is more permissive to active transcription (6–10). Additionally, re-positioning genes away from the nuclear periphery is associated with increased transcription (11–14).

At the sub-chromosomal level, chromatin is further organized into clusters of short-range interactions and chromatin loops. These intra-chromosomal interactions can result in specific long-range chromosome folding patterns that may be species- or context-specific (5). Together, this intricate 3D genome structure at both the whole- and sub-chromosome levels is essential for properly regulating gene expression in different developmental and tissue-specific contexts (15–19).

One prime example of how 3D chromosome architecture facilitates accurate gene expression is sex chromosome dosage compensation (DC). DC classically refers to the normalization of expression between sex chromosomes and autosomes, while sex chromosome dosage balance is the process of equalizing gene expression levels of sex-linked genes between homogametic (XX or ZZ) and heterogametic (XY or ZW) sexes (20, 21). Here, we refer to both of these two processes collectively as DC. Studies in diverse eukaryotic species have revealed an array of epigenetic mechanisms for establishing DC, all of which are associated with sex-specific alterations to the 3D structure of the sex chromosomes (22–28). For example, in mammals where X-linked expression is reduced in females (XX) by inactivation of a single chX, the inactive chX is more peripheral in the nucleus than the active chX, and the inactive chX adopts a unique, highly compact 3D configuration compared to the active chX (29).

The first models for the evolution of DC were proposed by Susumo Ohno. Ohno predicted that mechanisms would evolve to not only downregulate X-linked expression in the homogametic sex (XX) to maintain balanced chX expression between the sexes, but also to upregulate X-linked expression in both sexes to balance chX expression with autosomal expression (30). This two-step model for achieving DC is known as Ohno’s hypothesis, and it remains the prevailing model for the evolution of DC. Yet, studies both supporting and refuting particularly the latter part of Ohno’s hypothesis exist, leaving this model still widely debated (31–35).

Interestingly, the few existing studies in species with female heterogametry (ZW sex chromosomes) have suggested that DC mechanisms may be absent or incomplete in these lineages (36–44). However, previous expression-based studies demonstrated that Lepidopteran insects are an exception to this rule, where Z-linked gene expression in males (ZZ) is repressed to match the gene expression of the single chZ in females (ZW) (20, 45–50). In *Bombyx mori*, DC occurs in late embryonic development (45). Yet whether one Z is completely silenced (similar to X-inactivation in mammals) or both are partially repressed (similar to *C. elegans*) in Lepidopteran DC remains unclear. Furthermore, how 3D genome organization may more broadly be linked to transcriptional regulation in Lepidoptera is unknown, as the 3D organization of chromosomes in interphase has never been visualized in any moth or butterfly.

Here, we use Oligopaint DNA FISH to visualize autosomes and the Z sex chromosome (chZ) in male and female *B. mori* embryonic and larval cells. We reveal that *B. mori* chromosomes form distinct CTs, with both CT volume and position in the nucleus being influenced by both genomic size and gene density. Additionally, we show that gene-poor chromosomes harbor more robust long-range *cis* interactions than their gene-rich counterparts. We further show that after DC is established in late embryos, both copies of chZ in males are highly similar in volume, shape, and chromosome folding patterns. This finding suggests a mechanism in which both chZs in males are downregulated by half to achieve DC, similar to *C. elegans* (51). Furthermore, in support of Ohno’s hypothesis, we also find that the chZ in females is repositioned away from the nuclear edge when DC is established, and this central positioning of the chZ correlates with an increase in Z-linked gene expression by RNA sequencing (RNA-seq). This result is again similar to findings in *C. elegans* (52), further supporting a model in which DC in Lepidopteran insects and nematodes are established via similar, convergent mechanisms. Interestingly, both *C. elegans* and *B. mori* harbor holocentric chromosomes (where centromeres form all along the length of the chromosome), raising the possibility that the holocentric chromosome structure may influence the evolution of DC mechanisms. Together, these findings present the first visualization of 3D genome organization in any Lepidopteran species, illustrate that DC is achieved in *B. mori* by partially repressing both copies of the chZ in males, and reveal the first non-sequencing-based evidence for Ohno’s theory of the two-step mechanism of DC.

## Results and Discussion

### *B. mori* interphase chromosomes are highly compact

Before specifically interrogating chZ organization in *B. mori*, we needed to establish basic principles of interphase chromatin organization in this species. We began by measuring nuclear volume in *B. mori* embryonic cells using DAPI staining, which revealed that *B. mori* nuclei are extremely compact for their DNA content. The *B. mori* genome is over three times as large as the *D. melanogaster* genome (53, 54), and therefore, we expected *B. mori* nuclei to be three times as large. However, our measurements revealed instead that *B. mori* diploid nuclei are similar in volume to *Drosophila* diploid nuclei (Figure S1A). This finding suggests that extreme genome compaction may occur in moths relative to flies.

One possibility is that *B. mori* interphase CTs are more intermixed than fly CTs, allowing for a larger genome to occupy a smaller nuclear space. Supporting this hypothesis, the Condensin II complex that is required for CT separation in flies is reportedly incomplete in *B. mori* (5, 55). Thus, to measure CT intermixing, we labeled interphase chromosomes in *B. mori* embryonic nuclei with whole chromosome Oligopaints for six autosomes (chromosomes 4, 7, 15, 16, 17, and 23; Table 1) of the 28 *B. mori* chromosomes (56, 57) (Figure 1A). This assay clearly revealed the formation of spatially distinct CTs (Figure 1B), with the majority of cells (50-80%) harboring no overlap at all between any given representative CT pair (Figure 1C-1D). Also, contact between homologous chromosome copies occurs at comparable rates to heterologous contacts for two chromosomes of similar size (Figure 1C, 1E, S1B), supporting previous findings that somatic homolog pairing is absent in moths (55, 57). These results suggest that CT size is the main factor dictating CT-CT contacts in *B. mori*. However, size cannot fully explain all observed *trans* interactions, as ch7 has more heterologous and homologous contacts than ch16 (Figure 1C-1E), despite these two chromosomes being similar in size. Together, these assays demonstrate that *B. mori* harbor spatially distinct CTs and that high levels of CT intermixing cannot explain the small size of *B. mori* nuclei.

**Table 1.** Chromosome and chromosome paint information.

| Chrom. | chromosome size (bp) | size painted (bp) | paint start | paint stop | density (probes/kb) | total # of oligos | Gene density (genes per Mb) |
| --- | --- | --- | --- | --- | --- | --- | --- |
| 4 | 18737234 | 18639239 | 282 | 18639521 | 1.5 | 26841 | 39.71 |
| 7 | 13944894 | 13868845 | 35931 | 13904776 | 3 | 42625 | 36.00 |
| 15 | 18440292 | 18354755 | 21089 | 18375844 | 1 | 17756 | 43.60 |
| 16 | 14337292 | 14275583 | 27737 | 14303320 | 1.5 | 20190 | 43.66 |
| 17 | 16840672 | 16806551 | 9415 | 16815966 | 1.5 | 23834 | 38.30 |
| 23 | 21465692 | 21339065 | 123188 | 21462253 | 1.5 | 30506 | 35.82 |
| Z (1) | 20666287 | 20578020 | 37936 | 20615956 | 1.5 | 29784 | 32.18 |

**Figure 1.**
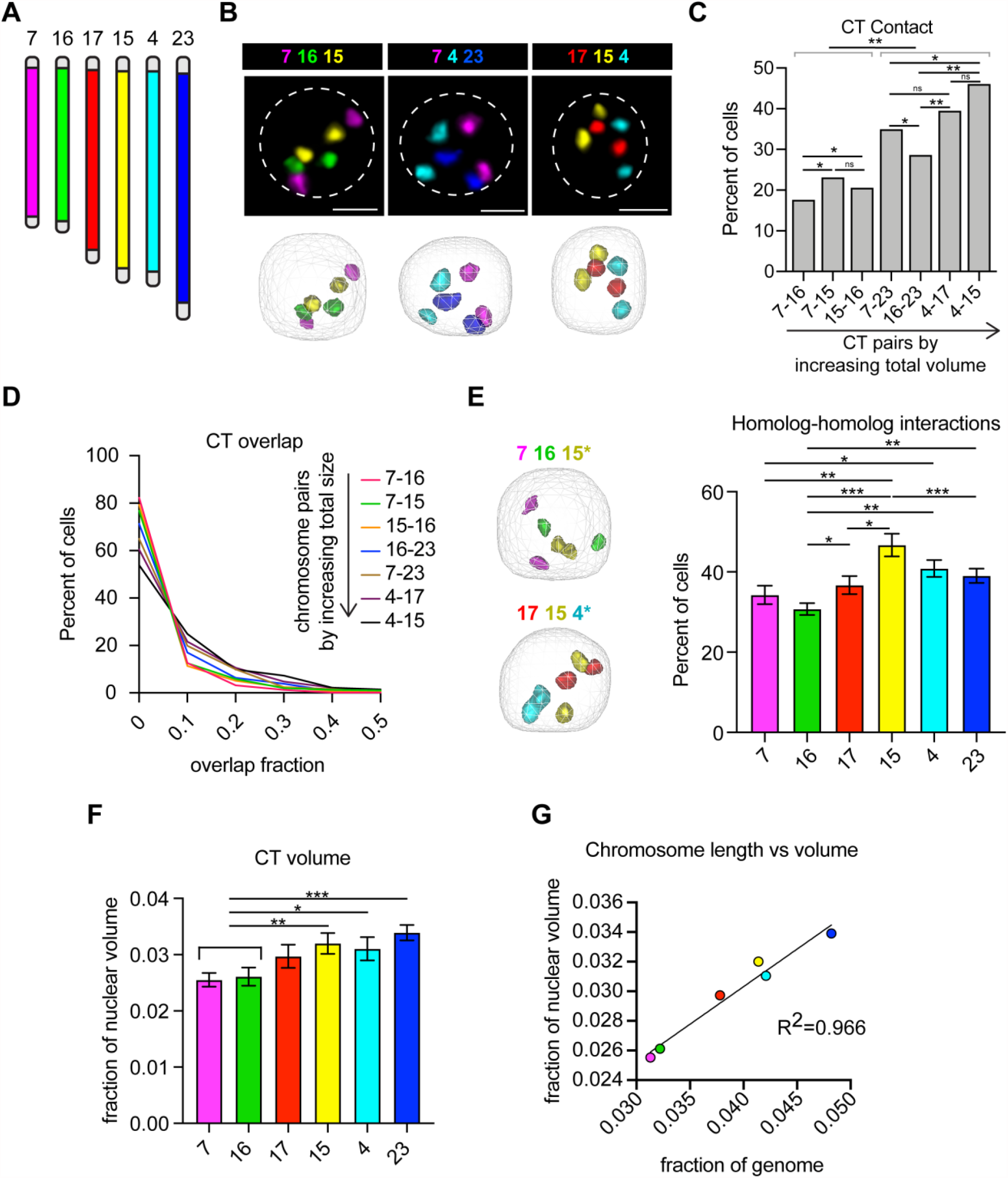
*B. mori* interphase chromosomes are compact and non-randomly organized. **A)** Schematic of whole chromosome Oligopaints used in Figure 1. Chromosomes and chromosome pairs are ordered by size (small to large) in all panels. **B)** Top: Representative Oligopaint DNA FISH images showing *B. mori* CTs. Scale bar = 2.5 µm. Dotted line indicates approximate nuclear edge. Bottom: 3D rendering of cells from TANGO. **C)** Bar graph showing the absolute contact frequency for all replicates (including at least 3 biological replicates per chromosome pair). Statistics: Fisher’s exact test comparing contact versus no contact. *p = 0.003 - 0.0003; **p <0.0003. **D)** Histogram showing CT overlap as a fraction of the smaller chromosome (for example, 15-16 overlap fraction is the fraction of ch16 colocalizing with ch15). All pairwise comparisons are statistically significant (Mann-Whitney Test, p<0.0001). **E)** Left: Representative 3D rendering of nuclei, as in B (bottom), showing homolog-homolog interactions of the chromosome pair indicated with an asterisk. Right: Quantification of homolog-homolog interactions shown as the fraction of cells harboring any voxel overlap between the two homologs of a single chromosome (indicated on the X-axis). Bars show the average between biological replicates. Error bars show standard error of the mean. Statistics = unpaired t-tests. *p<0.05, **p<0.01, ***P<0.001. **F)** Quantification of CT volume. Bars show average CT volume as fraction of nuclear volume across biological replicates. Statistics = unpaired t-tests. *p=0.035, **p=0.008, ***p=0.0006. **G)** Dot plot showing chromosome genomic length relative to total genome size (X-axis) versus CT volume relative to nuclear volume (Y-axis). R^2^=0.966. For all data shown in Figure 1, at least 3 biological replicates (embryos) were analyzed for each chromosome and at least 100 cells per embryo. Error bars show standard error of the mean.

Another possible explanation for the small size of *B. mori* nuclei is that chromatin within CTs may be more tightly packaged in *B. mori* than in flies, leading to smaller CTs and overall reduced nuclear volumes. We therefore measured CT volumes, which illustrated that on average, each chromosome occupies only 2-4% of the nucleus (Figure 1F). This amount is precisely the percent of the *B. mori* genome each chromosome encompasses, and thus, the genomic length of chromosomes scales nearly perfectly with CT volume (Figure 1G). Importantly, the volume of *B. mori* chromosomes is, indeed, significantly smaller than that of fly chromosomes of similar genomic size (Figure S1C). These findings suggest that *cis* compaction is highly robust in *B. mori*, but basic principles of large-scale nuclear organization such as CT formation are largely conserved in this species.

### *B. mori* autosomes are tightly folded

Our above findings suggest that sub-chromosomal compaction is more robust in moths than flies. As previous studies in flies illustrated that within CTs, chromosomes can have preferential folding configurations (5), we reasoned that one way to extensively compact chromosomes would be to fold them into tighter configurations. To test this hypothesis and to broadly assess long-range *cis* interactions in *B. mori* nuclei, we used Oligopaints to label ∼3 Mb stripes at both telomeres (tel1 and tel2) and in the center (mid) of ch7, ch15, and ch16 (Figure 2A-B and Table 2). Based on contact patterns between these stripes, eight possible folding configurations can be detected with this assay (Figure 2C; (5)). Among these configurations, the four in which the two telomeres are not in contact were classified as “unfolded” configurations, and the four in which the telomeres are in contact were classified as “folded” (Figure 2C). This assay revealed that in both embryonic and larval nuclei, *B. mori* chromosomes tend to adopt a single “folded” configuration where all three domains are in contact (Figure 2D-E). Ch7, which is small and gene-poor, is found in this completely folded configuration at even higher frequencies (∼75%). Importantly, this folding pattern is dramatically more compact than the folding patterns observed in fly cells, where only 30% of chromosomes are in a configuration with all stripe domains in contact (Figure S1D; (5)). Notably, for all three examined *B. mori* chromosomes, both homologous copies are folded similarly in 60-75% of nuclei, with the majority of these nuclei harboring two folded copies (Figure 2F-G), suggesting minimal intra-nuclear homolog heterogeneity. This result is in contrast to findings from human cells which showed that there are high levels of intra-nuclear homolog heterogeneity in interphase (58). This extensive level of folding may at least partially explain why *B. mori* nuclei are so much smaller than their fly counterparts. Together with the above studies, these data show for the first time basic principles of interphase nuclear organization in *B. mori*.

**Table 2.** Stripe sub-library paint information.

| Chrom. | Tel1 size (Mb) | Mid size (Mb) | Tel2 size (Mb) | density (probes/kb) |
| --- | --- | --- | --- | --- |
| 7 | 2.77 | 2.76 | 2.77 | 3 |
| 15 | 3.66 | 3.68 | 3.62 | 1 |
| 16 | 2.84 | 2.84 | 2.84 | 1.5 |
| 23 | 3.16 | 1.65 | 3.30 | 1.5 |
| Z (1) | 3.12 | 1.58 | 3.06 | 1.5 |

**Figure 2.**
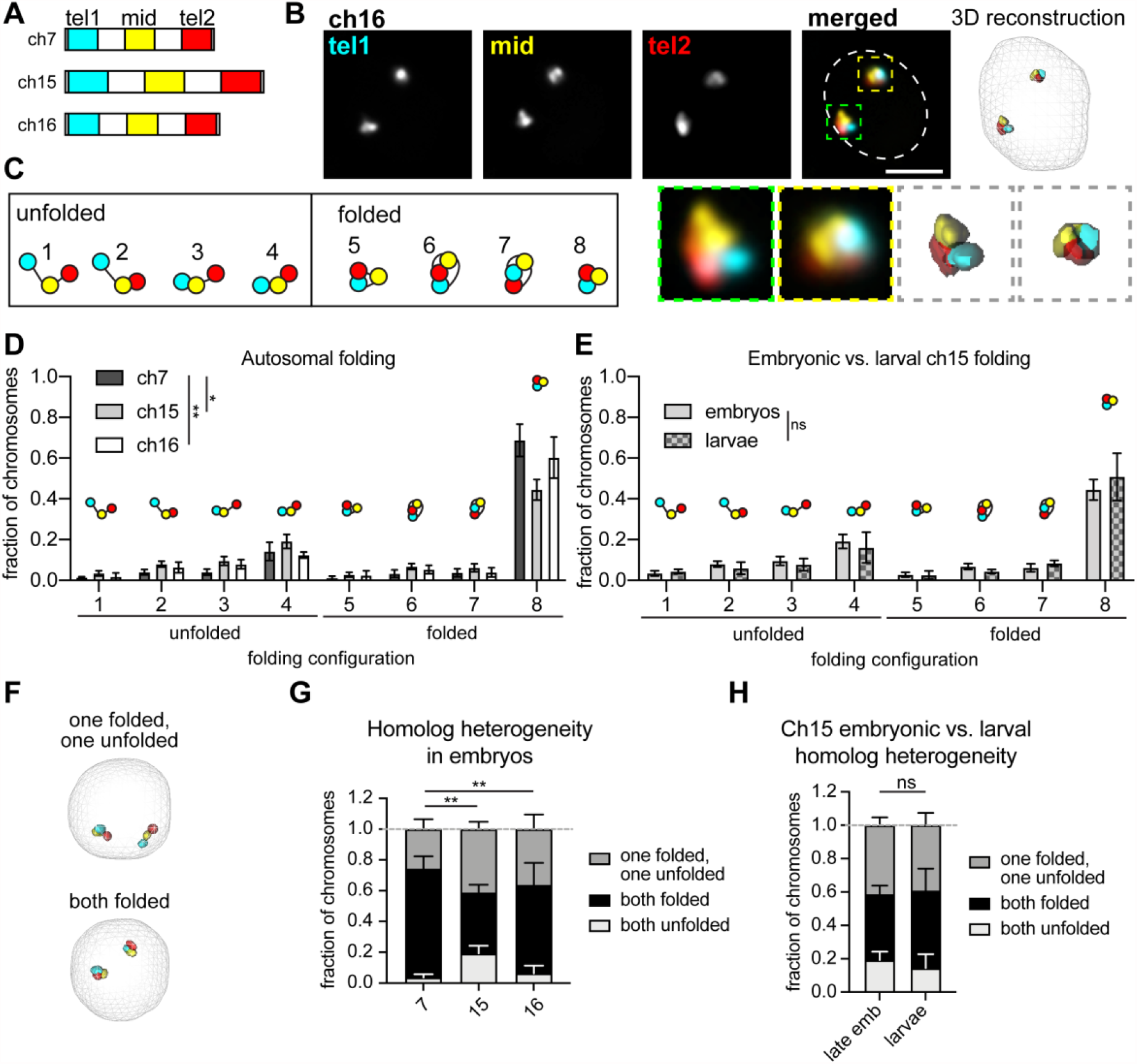
*B. mori* autosomes are tightly folded. **A)** Schematic of stripe Oligopaints used in Figure 2. Each chromosome is divided into 5 equally sized domains, and the first (tel1), middle (mid), and last (tel2) are labeled as indicated. **B)** Top left: Representative Oligopaint DNA FISH images showing stripe paints for ch16. Scale bar = 2.5 µm. Dotted line indicates approximate nuclear edge. Top right: 3D rendering of cell from TANGO. Bottom left: zoom of individual chromosomes labeled with Oligopaints from above. Bottom right: zoom of TANGO 3D rendering. **C)** Schematic of 8 possible folding configurations detected by this assay. Configurations with telomere-telomere contact (cyan and red) are categorized as “folded”, while configurations without telomere-telomere contact are classified as “unfolded”. **D)** Quantification of folding for ch7 (dark gray), ch15 (gray), ch16 (white) in embryos. Statistics = Chi-square test. *p = 0.01, **p = 0.003. **E)** Quantification of folding for ch15 in late embryos and larvae. Statistics = Chi-square test. **F)** Representative 3D rendering of nuclei from TANGO, as in B, showing a cell with homologs in different folding configurations (top) and a cell with homologs in the same configuration (bottom). **G)** Quantification of homolog heterogeneity in embryos. Bars show the average between biological replicates and error bars show the standard error of the mean. Statistics = Chi-square test. **p = 0.002 - 0.009. **H)** Quantification of homolog heterogeneity for ch15 in embryos and larvae. Statistics = Chi-square test. For all graphs, bars show the average of at least three biological replicates with at least 100 cells analyzed from each replicate. Error bars show standard error of the mean.

### ChZ is more compact in males than females

As DC in all studied species has been associated with sex-specific alterations to the 3D structure of the sex chromosomes, we examined the 3D organization of the *B. mori* chZ sex chromosome by FISH to interrogate possible structural changes associated with dosage compensation (DC). In *B. mori*, a transcriptome study found that Z-linked gene expression is reduced in males (ZZ) to match the expression of the single chZ in females (ZW), and this equalization occurs by late embryonic development (45). However, whether one chZ is completely silenced (as in mammals) or both are partially and equally repressed (as in *C. elegans*) in moths DC is unknown. In agreement with previous studies in Lepidoptera (59), we saw no cytological evidence of Barr body formation in male (ZZ) nuclei, suggesting complete Z-inactivation may not occur (Figure S2A). However, we noted that nuclei from both males and females appear to be completely devoid of dense heterochromatin based on a lack of DAPI-bright signal (Figure S2A). In agreement with this observation, previous ChIP-seq studies revealed only minimal H3K9me2/3 genome-wide in *B. mori* cultured cells (60). Therefore, we reasoned that it is possible that complete Z-inactivation could occur even in the absence of a cytological Barr body.

Irrespective of Barr body formation, we reasoned that if Z-inactivation occurs in *B. mori*, one male chZ homolog would be significantly more compact than the other by DNA FISH. Visualizing chZ by FISH would also reveal if, instead, both chZ homologs in males are similar in size but are more compact than the single chZ in females, indicating partial repression of both chZ copies in males. To this end, we used Oligopaints for the *B. mori* chZ that were similar in design to those used to label autosomes (57). As a control, we used ch23, which is a similar genomic size and gene density to chZ (Figure 3A and Table 1). We examined chZ organization at three distinct developmental timepoints: a mid-embryonic stage (before DC is achieved), a late embryonic stage (when DC is achieved), and a late larval stage (5^th^ instar, at which DC should be stable)(45). At each of these time points, we measured CT volume and position in the nucleus to look for differences between the male and female chZ relative to ch23.

**Figure 3.**
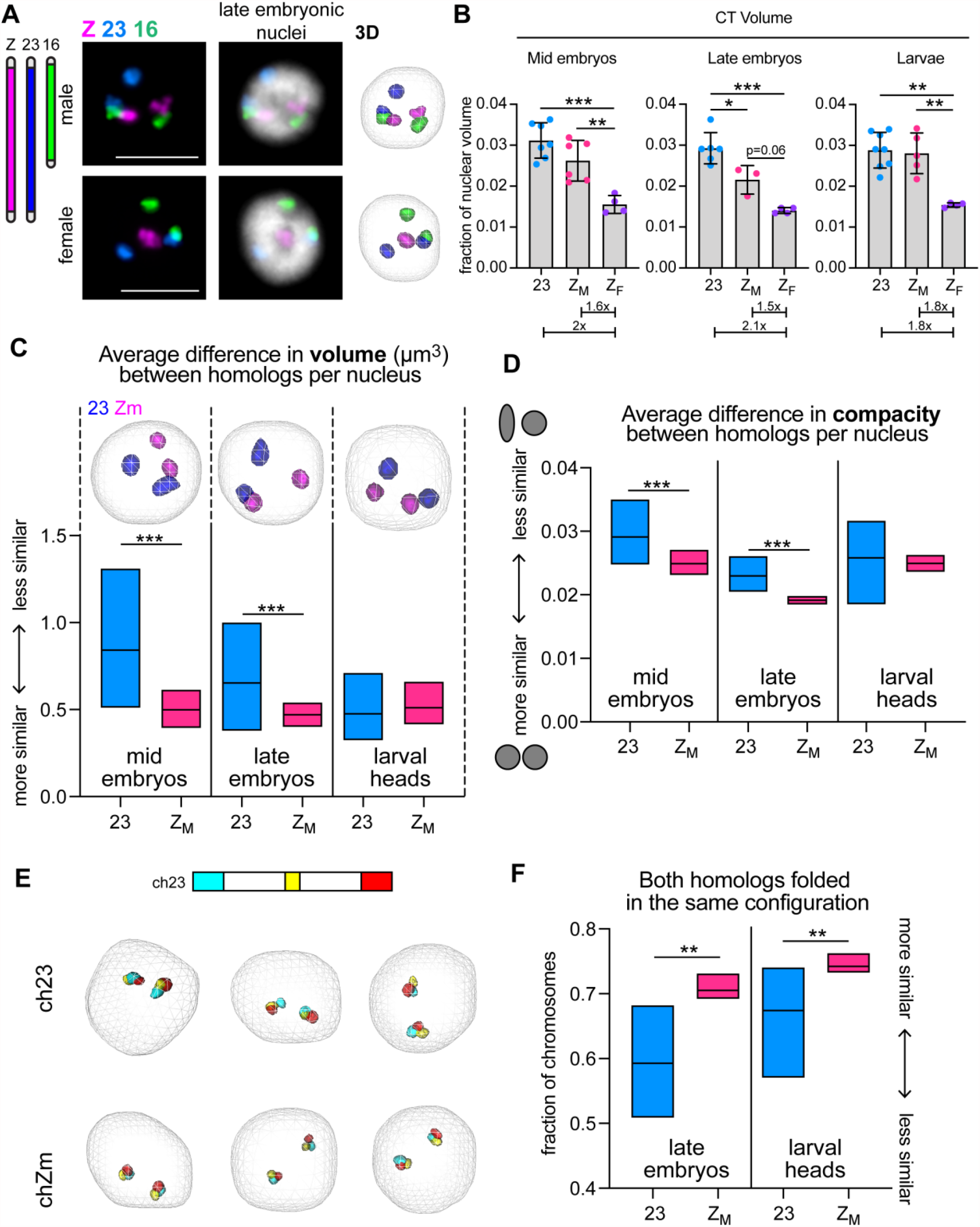
Male Z chromosomes are similar in size and shape, suggesting partial repression of both homologs. **A)** Left: Schematic of whole chromosome Oligopaints used in these experiments. Right: Representative Oligopaint DNA FISH images showing *B. mori* CTs for chZ, 16, and 23. Nucleus from male late embryo (ZZ, top) and nucleus from female late embryo (ZW, bottom) is shown. Scale bar = 5 µm. Right: 3D rendering of cells from TANGO. **B)** Quantification of total CT volume per nucleus at different developmental stages. Each dot represents average of biological replicates (embryo or larva). Bars show average between biological replicates and error bars show standard deviation. Below, average ch23 or chZ_M_ value is normalized to average Z_F_ value. For all data in A-B, at least 100 cells were analyzed per replicate. Error bars show standard error of the mean. Z_M_ = chZ in male nuclei; Z_F_ = chZ in female nuclei. Statistics = unpaired t-tests. *p = 0.01 - 0.05, **p = 0.001 - 0.005, ***p < 0.001. **C)** Top: 3D rendering of male nuclei from TANGO showing ch23 (blue) or chZ (pink) at indicated developmental time points. Bottom: Quantification of interhomolog differences in volume (µm^3^) for ch23 (blue) or chZ (pink) in male embryos or larvae. Mid-line = mean. Statistics = Mann-Whitney Test. *** p < 0.0001. **D)** Quantification of interhomolog differences in shape as measured by compacity (ratio between volume and surface area of object, where 1 = perfect sphere) for ch23 (blue) or chZ (pink) in male embryos or larvae. Mid-line = mean. Statistics = Mann-Whitney Test. ***p < 0.0001. **E)** Top: Schematic of stripe Oligopaints for ch23 and chZ. 3 Mb domains are painted at both telomeres (cyan, stripe 1; red, stripe 3) and a 1.5 Mb domain is painted in the middle of the chromosome (yellow, stripe 2). Middle and Bottom: representative 3D renderings from TANGO of male nuclei with stripe paints for ch23 (middle) and chZ (bottom). **F)** Quantification of the fraction of cells with both homologs for either ch23 (blue) or chZ (pink) folded in the same configuration based on stripe paint assay. Mid-line = mean. Statistics = unpaired t-test. Late embryos **p = 0.003; larval heads **p = 0.005.

CT volume measurements before and after the establishment of DC revealed that total chZ volume per nucleus in males is less than that of total ch23 volume, with this difference being statistically significant after DC is established. After DC in late male embryos, both chZ copies together occupy only 1.8% - 2.4% of the nuclear volume, while both ch23 copies occupy 2.9% - 3.8% (Figure 3B, S2B, S2C). This reduced volume of chZ compared to ch23 suggests that chZ is more compact than ch23 in males. Additionally, both before and after DC in embryos, the total volume of two chZ in males is only 1.6x greater (not 2x greater) than the single chZ volume in females, suggesting that the chZ in females is not as compact as the chZs in males (Figure 3B). This result is highly reminiscent of FISH-based assays in *C. elegans* showing that each chX in hermaphrodite worms is more compact than the single chX in males (25, 28).

Interestingly, by the 5^th^ instar larval stage, we found that the total volume for both copies of chZ and ch23 in males are equal, and the volume of the single chZ in females is half this volume (Figure 3B, right), indicating that similar levels of compaction are occurring for chZ and autosomes in both males and females by this developmental stage (when DC should be stable (45)). This seeming loss of hypercompaction of chZ in males in 5^th^ instar larvae suggests that perhaps an alternative, compaction-independent DC mechanism operates during this developmental window. Another possibility is that global changes in gene expression profiles in preparation for metamorphosis may result in global changes in chromosome compaction independent of DC in 5^th^ instar larvae (61).

### Both male chZ homologs are similar in size, shape, and folding configuration

While our measurements of total chZ volume per nucleus support the repression of Z-linked genes in male late embryos, whether both chZ copies are partially downregulated or one chZ is completely inactivated in males is still unclear. To address this specific question, we measured the differences in volume and shape between the two homologous chZ copies and the two ch23 copies within individual male nuclei (defined here as intra-nuclear difference). If both male chZs are partially and equally repressed, we would expect both chZ copies to be similar in volume and shape. However, if one chZ is inactivated, the two chZ copies would be different in volume and shape.

Measuring the intra-nuclear volume difference for chZ and ch23 revealed that in both mid and late embryos, the two chZ copies in males are significantly more similar in volume than the two ch23 copies (Figure 3C and Figure S2D), suggesting that the two chZ copies are similarly compacted. In agreement with this finding, measuring the intra-nuclear difference in shape illustrated that both chZs are significantly more similar in shape compared to ch23 at both embryonic stages (Figure 3D and Figure S2E). Curiously, these differences between chZ and ch23 are again gone by the larval stage (Figure 3C-D and Figure S2D-E), supporting the idea that a compaction-independent mechanism facilitates DC at this developmental stage. Furthermore, at all time points, ch23 shows much higher variance for intra-nuclear size and shape than chZ, indicating not only higher homolog-homolog variability, but higher cell-to-cell variability for ch23. Similarly high variance was observed for ch16 (Figure S2F), suggesting that this variability may be exhibited by all autosomes. The finding that the two chZs in males are highly similar in volume and shape supports a model of DC in which expression from both chZs is partially and equally reduced.

To further investigate chZ compaction and organization, we examined the folding configuration of chZ and ch23 by measuring contact patterns between sub-chromosomal stripes, as previously described for autosomes (Figure 2). This assay revealed that on average in a population of cells, the chZs in males are more often completely folded (with contact between all three stripes) compared to ch23 (Figure 3E and Figure S2G). Next, we measured intra-nuclear folding heterogeneity by quantifying how often the two homologs in the same nucleus are either both unfolded or both folded for chZ and ch23. We reasoned that if both chZ copies are equally repressed, the two homologs would harbor similar folding patterns in most nuclei, whereas if one chZ is inactivated, there would be higher intra-nuclear heterogeneity. Indeed, we found that chZ homologs in males are significantly more often in the same configuration compared to ch23, and these results are similar during both timepoints after DC is established (Figure 3F).

Taken together, these data suggest that Z-linked repression in *B. mori* males is achieved by partially repressing both chZ copies and not by entirely inactivating one copy. This mechanism is highly reminiscent of DC in *C. elegans*, despite these two lineages bearing evolutionarily distinct ZW and XY sex chromosomes. Intriguingly, like *B. mori* and all Lepidoptera, *C. elegans* harbor holocentric chromosomes in somatic cells. Whether or not this holocentric configuration inhibits the inactivation of an entire chromosome, or perhaps facilitates partial repression of both sex chromosomes in the homogametic sex, remains to be explored.

### Changes in female chZ volume and nuclear position correlate with changes in Z-linked expression

We next wanted to determine whether there are any differences in nuclear position of chZ in males versus females. To address this question, we measured the position of autosomes and chZ in the nucleus using a shell analysis. In this assay, nuclei are divided into nine concentric shells of equal volume and the total paint volume in each shell is measured (Figure 4A). Shell analysis of autosomes revealed that gene-poor chromosomes (like ch23) are preferentially located at the nuclear periphery in late embryos (Figure S3A). This is in agreement with data on CT position in other species (3–5) and in agreement with the fact that chromatin at the nuclear periphery is broadly repressed (6–10). Interestingly, we also observed that small chromosomes are at the nuclear periphery more often than larger chromosomes, even when they are gene-rich (Figure S3A). This finding is distinct from what has been previously shown in humans, where smaller chromosomes tend to be more central in the nucleus (2).

**Figure 4.**
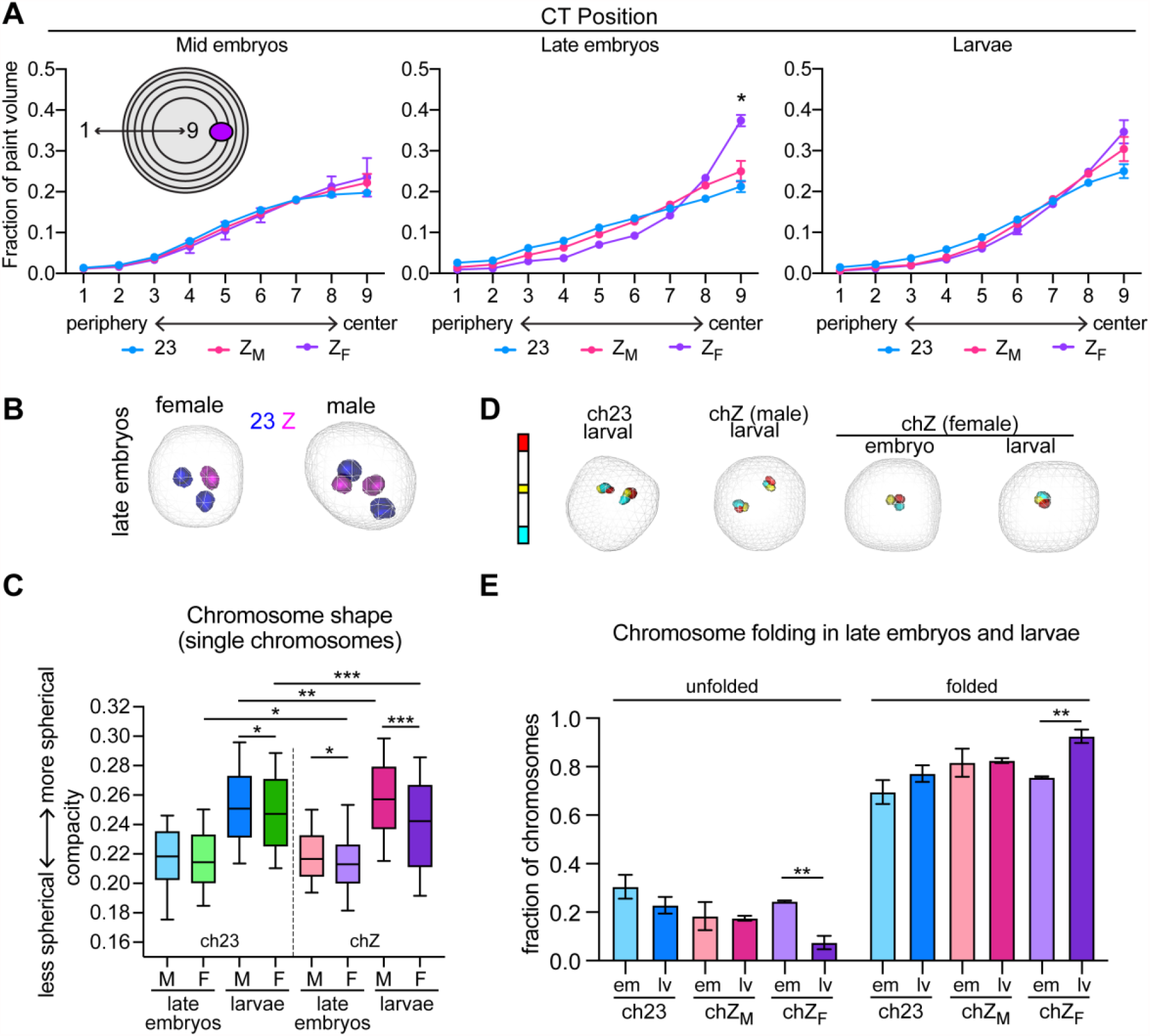
Changes in female Z chromosome volume and nuclear position correlate with changes in Z-linked gene expression. **A)** Quantification of CT position. Nuclei are divided into 9 shells of equal volume, with shell 1 being most peripheral and shell 9 being most central. The 3D volume of each chromosome paint in each nuclear shell is quantified. Shown are averages and standard error across at least three biological replicates (embryos or larvae), where at least 100 cells were measured per replicate. Statistics = Chi-square test. *p = 0.03. **B)** 3D rendering of female (left) or male (right) nuclei from TANGO showing ch23 (blue) or chZ (pink) in late embryos. **C)** Tukey box and whisker plot showing quantification of chromosome shape (compacity; Y-axis) for individual chromosomes at the indicated time points. Bold midline = median; dashed lines indicate quartiles. Statistics = Mann-Whitney Tests. *p = 0.01 - 0.05, **p = 0.001 - 0.005, ***p < 0.001. **D)** Left: schematic of stripe Oligopaints for ch23 and chZ. Right: 3D rendering of stripe paints in larval or embryonic male or female individuals, as indicated. **E)** Quantification of fraction of ch23 (blue), chZ_M_ (pink), or chZ_F_ (purple) in unfolded (left) or folded (right) configuration in late embryos (em) and larvae (lv). Statistics = Fisher’s exact test comparing unfolded versus folded chromosomes in embryos and larvae. **p = 0.0046.

We next tested whether chZ is positioned similarly to ch23, which is similar in both size and gene density, and found no significant differences in their nuclear positions in males at any developmental stage (Figure 4A, 4B and Figure S3B). Furthermore, we calculated the intra-nuclear difference in position for the two chZ and ch23 copies in males by measuring the distance of individual homologs from the nuclear periphery. Again, no significant difference was observed between chZ and ch23 (Figure S3C). Strikingly, in females, we observed a substantial difference in the nuclear position of chZ relative to ch23. We found that in late embryos, when DC is established (45), the female chZ is repositioned to the center of the nucleus (Figure 4A-B and Figure S3B). This shift toward the nuclear center persists into the larval stage but to a lesser extent. Notably, the single chX in *C. elegans* males is also preferentially positioned in the nucleus after DC, although in this case the male chX is positioned close to the nuclear periphery and interacts with nuclear pore proteins (28, 62).

We wondered whether this shift in position of female chZ to the center of the nucleus corresponded to possible decompaction and increased gene expression, since the nuclear interior is more permissive to active transcription. Additionally, an increase in Z-linked expression in females would directly support Ohno’s hypothesis (30). To test whether changes in chromosome compaction occur at this developmental time, we measured the shape of chZ and ch23 in males and females. Our earlier measurements revealed no change in volume for chZ in females at the late embryonic stage, but differences in chromosome shape can alternatively indicate altered chromatin compaction, as chromosomes become more linear and less spherical when they decondense (5). As a reference, we verified that ch23 shape is similar between males and females throughout development (Figure 4B and 4C). When measuring chZ shape, we found that female chZ is significantly less spherical (and thus less compact) than both ch23 and male chZs after DC onset in late embryos and larvae (Figure 4C). This result is consistent with possible upregulation of female chZ during this time. Importantly, we also observed that male chZs are significantly more spherical (and thus more compact) than ch23 in larvae, consistent with persistent repressed expression at this later developmental stage.

To further interrogate whether female chZ decondenses at DC onset, we measured chromosome folding using stripe paints for chZ and ch23 in females and males. When we measured the folding of chZ in females at two post-DC time points, we found that chZ in late female embryos is significantly more often unfolded compared to chZ in larval females (Figure 4D, 4E). This unfolded configuration of chZ in females is consistent with decompaction and elongation of the female chZ at this stage and consistent with the significant shift in chZ nuclear position observed in late female embryos. Interestingly, we found that in males, chZ homologs are only slightly more often folded compared to ch23 homologs (Figure 4D, 4E). Together with our earlier results, these findings suggest that compaction and decompaction of male and female chZs, respectively, are modulated by different molecular mechanisms. The single female chZ unfolds to decondense, indicating changes in long-range *cis* interactions. In contrast, male chZs reduce in volume but undergo only a mild increase in long-range folding, suggesting mainly changes in local, short-range interactions. These distinct mechanisms of sex chromosome regulation lead to sex-specific alterations to 3D chromosome structure.

Finally, we sought to verify that the repositioning and decompaction of chZ in late female embryos corresponds to changes in Z-linked expression. We therefore re-analyzed existing *B. mori* sex-specific RNA-seq data from mid-embryos (before DC), late embryos (DC established), and larvae (DC stable) (45). Previously, these data were used to argue that the male chZ is repressed to establish DC by the late embryonic stage (45). Here, we wanted to focus specifically on the female chZ and investigate whether female Z-linked expression is upregulated during DC. Our new analyses of these data revealed that, indeed, there is a brief window of increased Z-linked expression in both females and males in late embryos, concurrent with a change in position of female chZ toward the nuclear interior (Figure S3D). Importantly, while Z-linked transcription is globally increased relative to the autosomal average (Figure S3D), female Z-linked expression at this time point is significantly enriched compared to male Z-linked expression, despite females having half the allelic copy number (Figure S3E). This hyperactivation of sex chromosomes followed by stabilized repression in the homogametic sex directly supports Ohno’s hypothesis of the evolution of DC (30), and our FISH-based studies are the first to link this increased transcription to changes in 3D position and organization of sex chromosomes in Lepidoptera. All of our findings are highly reminiscent of DC mechanisms in *C. elegans*, despite *C. elegans* being XY-bearing and *B. mori* being ZW-bearing. Whether or not the holocentric chromosome structure influences dosage compensation patterns remains to be explored.

## Conclusions

In this study, we use Oligopaints to visualize both autosomes and chZ in female and male *B. mori* at different stages of development before and after DC establishment. We show that *B. mori* nuclei are highly compact overall and that chromosomes are spatially partitioned into distinct CTs. These CTs occupy preferential positions in the nucleus that follow both conserved and divergent rules of organization, with small and gene-poor chromosomes being most peripheral. We show for the first time that both chZ copies in male silkworms are similarly condensed and folded, supporting a model where Z-linked expression is achieved by partially and equally repressing both chZ copies. Additionally, we find concomitant repositioning, decompaction, and transcriptional upregulation of chZ in females, directly supporting Ohno’s hypothesis for the evolution of DC (30). This work represents the first non-sequencing based evidence in support of this long-standing model. Furthermore, these studies revealed striking similarities between dosage compensation in moths and *C. elegans*, despite these two lineages harboring evolutionarily distinct sex chromosomes. Whether or not the holocentric chromosome structure can explain this convergent evolution remains unclear. Finally, while dosage compensation studies in other holocentric species have suggested that histone modifications (59) or specialized Condensin complexes (63) can regulate sex chromosome-linked expression, the molecular mechanisms regulating DC and overall nuclear organization in *B. mori* remain subjects for future interrogation.

## Acknowledgements

We would like to thank the members of the Lei and Drinnenberg labs, as well as Jamie Walters, Brian Oliver, and Jeannie Lee for critical reading of the manuscript and helpful discussion. We would also like to thank members of the Joyce lab for helpful discussion, and Dahong Chen for technical assistance with *Drosophila* neuron isolation.

## Methods

### *B. mori* strains

Moth embryos were obtained from Carolina Biological, Coastal Silkworms, Mulberry Farms or were freshly laid in the lab from moths derived from embryos from these sources. Some larvae were obtained from Rainbow Mealworms. Embryos were kept in diapause at 4°C for 3 mo to 1 y. For rearing, embryos were transferred to 28°C, and larvae were fed fresh mulberry leaves or powdered mulberry chow (Carolina Biological).

### *B. mori* staging, embryonic and larval cell slide preparation, and sexing

*B. mori* embryonic and larval stages were defines as follows: mid embryos = during embryonic diapause (at 4°C less than 1 mo); late embryos = post-diapause (at RT 7-10 days after removal from 4°C, before hatching); larvae = 5^th^ instar (approximately 3 in long and after cessation of eating).

For slide preparation, embryos were partially dechorionated in 50% bleach for 15 min at RT, followed by manual removal of the chorion and vitelline membrane with forceps in a glass dissecting dish containing RT Sf-900 media (Gibco). For mid embryos, cells were then harvested directly from the dissecting dish and settled on poly-L-lysine-treated glass slides for 30 min in media before fixing with 4% PFA in PBS. For late embryos, pre-larvae were manually dissected away from extra-embryonic tissue and cells were dissociated with papain and collagenase before settling on slides and fixing as above. For larval samples, heads from 5^th^ instar larvae were collected by decapitation, and cells were dissociated using Papain and Collagenase. Single embryos or single larval heads were used for slides involving Z chromosome analyses, and the number of Z chromosomes was used to sex the embryos. Larvae were sexed during dissection based on gonad morphology.

### *Drosophila* nuclear analyses and Oligopaint data

Images used for analyses of CTs and folding of *Bombyx* versus *Drosophila* chromosomes were from previously published studies (5, 64). For nuclei analysis in pupal neurons, pupal brains were dissected and neurons were isolated by FACS as previously described (65). Neurons were then settled on Poly-L lysine coated slides for 30 min in Schneider’s Media, fixed with 4% PFA for 10 min, washed with PBS, and stained with DAPI DNA stain.

### *B. mori* Oligopaint design and synthesis

Oligopaint libraries were previously published (57) and designed as previously described using the Oligominer pipeline (66). Coordinates and chromosome information for all paints can be found in Tables 1 and 2. For stripe paints, whole chromosome paints were multiplexed as previously described (57, 67) to allow for amplification sub-chromosomal stripes. Chromosomes 7, 15, and 16 contain 5 sub-chromosomal stripes approximately 3 Mb each. Of these, stripes 1, 3, and 5 were labeled to generate the 3-stripe pattern (tel1, mid, and tel2, respectively). Chromosomes Z and 23 contain 13 sub-chromosomal stripes approximately 1.5 Mb each. For chZ and ch23, stripes 1,2 (combined = tel1), 7 (mid), and 12, 13 (combined = tel2) were labeled to generate the 3-stripe pattern.

### Oligopaint DNA FISH in *B. mori* cells

Oligopaint synthesis and FISH were performed as previously described (5, 68). Briefly, for FISH, after slide preparation and fixation, slides were washed 3x 5 min each in 0.1% Triton X-100 in PBS (PBS-T^0.1%^) at RT, then permeabilized with PBS-T^0.5%^ for 15 min at RT. Cells were subsequently pre-denatured using the following washes: 1x 5 min in 2xSSCT (0.3 M NaCl, 0.03 M sodium citrate, 0.1% Tween-20) at RT, 1x 5 min 2xSSCT/50% formamide at RT, 1x 2.5 min in 2×SSCT/50% formamide at 92°C, 1x 20 min at 60°C in 2xSSCT/50% formamide. Primary Oligopaint probes in hybridization buffer (10% dextran sulfate/2xSSCT/50% formamide/4% polyvinylsulfonic acid) were then added to slides, covered with 22×22 mm cover glass, and sealed before being denatured at 92°C for 2.5 min. Slides were then placed in a humidified chamber at 37°C and incubated 16-18 h. The next day, slides were washed as follows: 2xSSCT at 60°C for 15 min, 2xSSCT at RT for 15 min, 0.2xSSC at RT for 10 min. Secondary probes (10 pmol/25 µL) containing fluorophores were added to slides, resuspended in hybridization buffer as described above, and covered with 22×22 mm cover glass and sealed. Slides were incubated at 37°C in a humidified chamber for 2 h before repeating the above washes. Finally, all slides were stained with DAPI DNA stain in PBS for 5 min, followed by 2x 5 min washes in PBS-T^0.1%^ before mounting in Prolong Diamond (Invitrogen). Slides were left to cure overnight at RT before sealing with nail polish.

### Imaging, quantification, and data analysis

Images were acquired on a Leica DMi6000 widefield fluorescence microscope using an HCX PL APO 63x/1.40-0.60 Oil objective or HCX PL APO 100x/1.40-0.70 Oil objective (Leica), and Leica DFC9000 sCMOS Monochrome Camera. DAPI, CY3, CY5, and FITC filter cubes were used for image acquisition. All images were processed using the LasX software and Huygens deconvolution software, and tiffs were created in ImageJ. For quantification, deconvolved images were segmented and measured using a modified version of the TANGO 3D-segmentation plug-in for ImageJ (69), using either the ‘Hysteresis’ or ‘Spot Detector 3D’ algorithms. Statistical analyses were performed using Prism 9 software by GraphPad. Figures were assembled in Adobe Illustrator.

### RNA-seq data and quantification

Raw RNA-seq data were downloaded from BioProjectID: PRJNA388026. Adaptors were detected by BBtools v38.82 (70). Low quality bases and adaptors were trimmed using Trimmomatic v0.39 (71) with the previously described parameters (45). The filtered paired-end reads were mapped to the Ensembl (2013) Bombyx mori (ASM15162v1) reference genome using bowtie v2.4.1 (72) with default parameters. SAM/BAM conversions, sorting, indexing and filtering were performed with SAMtools v. 1.10 (73). Aligned reads in the bam files were counted using subread featureCounts v2.0.1 (74) with default parameters. The GTF file was downloaded from Ensembl Bombyx_mori.ASM15162v1.48. To find the corresponding chromosome number for each gene, we downloaded the annotation file BombyxMoriComprehensiveGeneList.xlsx from https://sgp.dna.affrc.go.jp/ComprehensiveGeneSet/ (75). Raw read counts were normalized by FPRM (Fragments Per Kilobase Million).

All annotated genes were grouped into autosomal (A)- and Z-linked (Z) genes. To compute the log2(FPKM) values, we filtered out genes with raw counts equal to 0 in any sample at 78 h and 120 h embryos and larval heads. After filtering, the total gene number was 7210. Mean FPKM values of Z and all A were used to compute the Z:A ratios. Finally, we divided the ordered log2(FPKM) value of Z-linked gene expression into four quantiles from Q1 (low) to Q4 (high) as previously described (45). Welch Two Sample t-tests (two-sided) were used to compare the mean of Z-linked gene expression for male and female embryos at 78 h (mid), 120 h (late), and head (larval head) stages based on the log2(FPKM) value.

## Author Contributions

Conceptualization: L.F.R.; Oligopaints: L.F.R.; Image analysis: L.F.R.; RNA-seq analysis: Y.C.; L.F.R.; Writing - original draft: L.F.R.; Supervision: E.P.L.; Funding acquisition: E.P.L.

## Supplemental Figures and Legends

**Figure S1.**
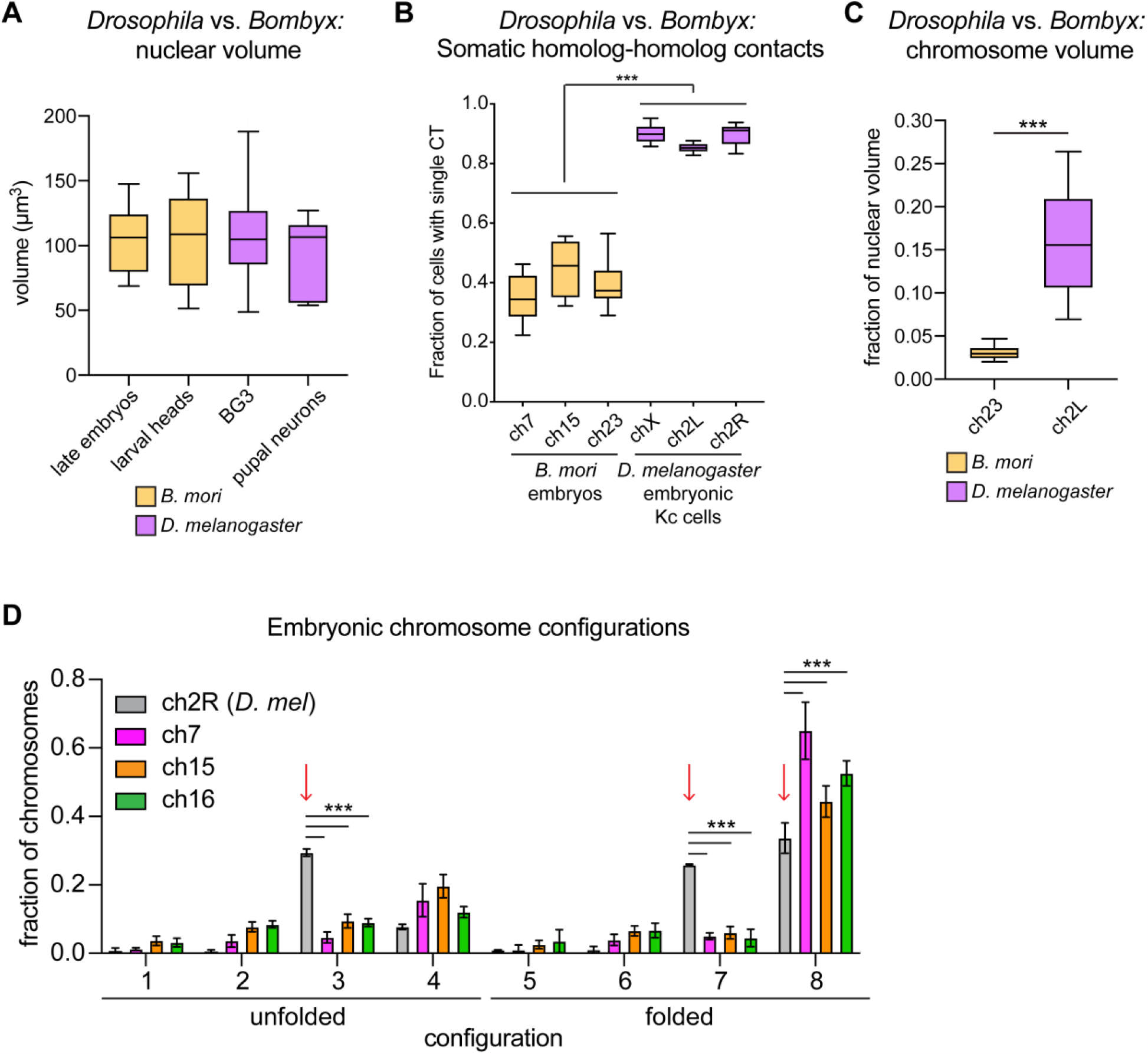
*B. mori* nuclei and chromosomes are relatively more compact than their *D. melanogaster* counterparts. A) Tukey box and whiskers plot showing the average nuclear volume of *B. mori* diploid cells from late embryos and larval heads (orange) and *D. melanogaster* BG3 cultured cells and pupal neurons (purple). Plots show the averages from 7-15 biological replicates. B) Tukey box and whiskers plot showing the average fraction of cells harboring a single contiguous CT for three representative chromosomes in *B. mori* embryos (orange) and 3 representative chromosome arms in *D. melanogaster* embryonic Kc cells (purple). ***p<0.0001; Mann-Whitney Test for all pairwise comparisons. C) Tukey box and whiskers plot showing the average size (fraction of nuclear volume) for *B. mori* chromosome 23 (orange) from embryonic cells, and *D. melanogaster* ch2L (purple) from BG3 cells. Data shown represent a single biological replicate (n = 1500 cells from *B. mori* and n = 2017 cells from *D. melanogaster*), but similar results were found in 3-5 additional replicates. ***p<0.0001; Mann-Whitney Test. D) Quantification of folding for *B. mori* embryonic ch7 (magenta), 15 (orange), and 16 (green), and *D. melanogaster* ch2R from embryonic Kc cells (gray). Red arrows indicate configurations enriched in Kc cells (3, 7, 8). Bars show the average of at least three biological replicates with at least 100 cells analyzed from each replicate. Error bars show standard deviation. ***p<0.0001; Multiple t-tests.

**Figure S2.**
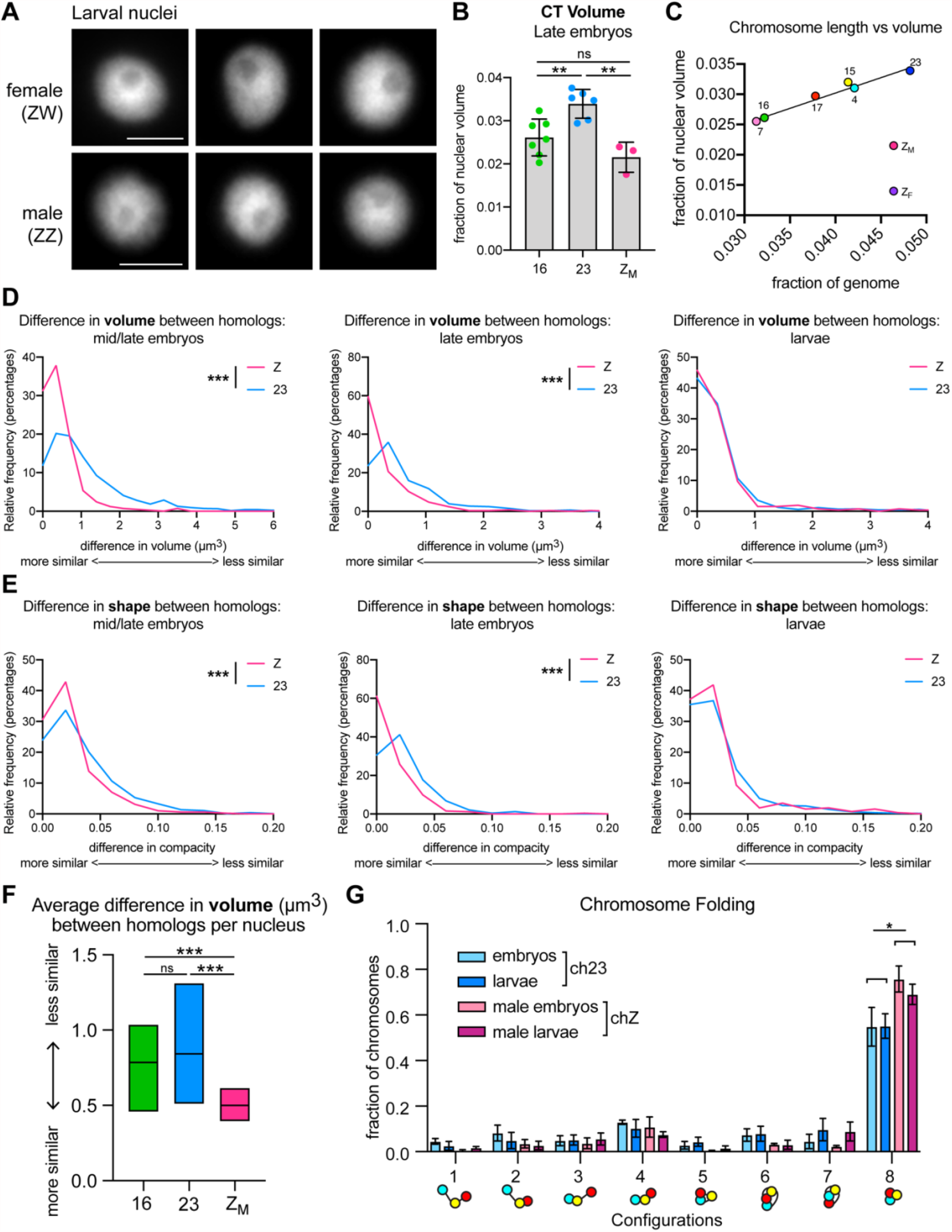
Both male chZ homologs are more similar in size and shape than the ch23 homologs. A) DAPI staining of representative larval nuclei from B. mori females (top) and males (bottom). Scale bar = 2.5 µm. B) Quantification of CT volume in late embryos. Each dot represents the average of a biological replicate. Bars show average CT volume as a fraction of nuclear volume between replicates. Error bars show standard error of the mean. Z_M_ = chZ in male nuclei. Statistics = unpaired t-tests. **p = 0.001 - 0.005. C) Dot plot showing chromosome genomic length as a fraction of total genome size (X-axis) versus CT volume as a fraction of nuclear volume (Y-axis). R^2^=0.966 excluding chZ. Data shown are from late embryos. D) Frequency histograms showing the difference in volume (µm^3^) between homologs for chZ (pink) and ch23 (blue) in male mid/late embryos (left), late embryos (middle), and larvae (right). Data represent a single biological replicate. ***p<0.0001; Mann-Whitney Test. E) Frequency histograms showing the difference in shape (compacity) between homologs for chZ (pink) and ch23 (blue) in male mid/late embryos (left), late embryos (middle), and larvae (right). Data represent a single biological replicate. ***p<0.0001; Mann-Whitney Test. F) Quantification of interhomolog differences in volume (µm^3^) in for ch16 (green), ch23 (blue), and chZ (pink) in male mid/late embryos. Mid-line = mean. Statistics = Mann-Whitney Test. *** p < 0.0001. G) Quantification of chromosome folding for ch23 (blue) and chZ (pink) in embryos and larvae. Bars show the average of at least three biological replicates with at least 100 cells analyzed from each replicate. Error bars show standard deviation. *p=0.01; Multiple t-tests.

**Figure S3.**
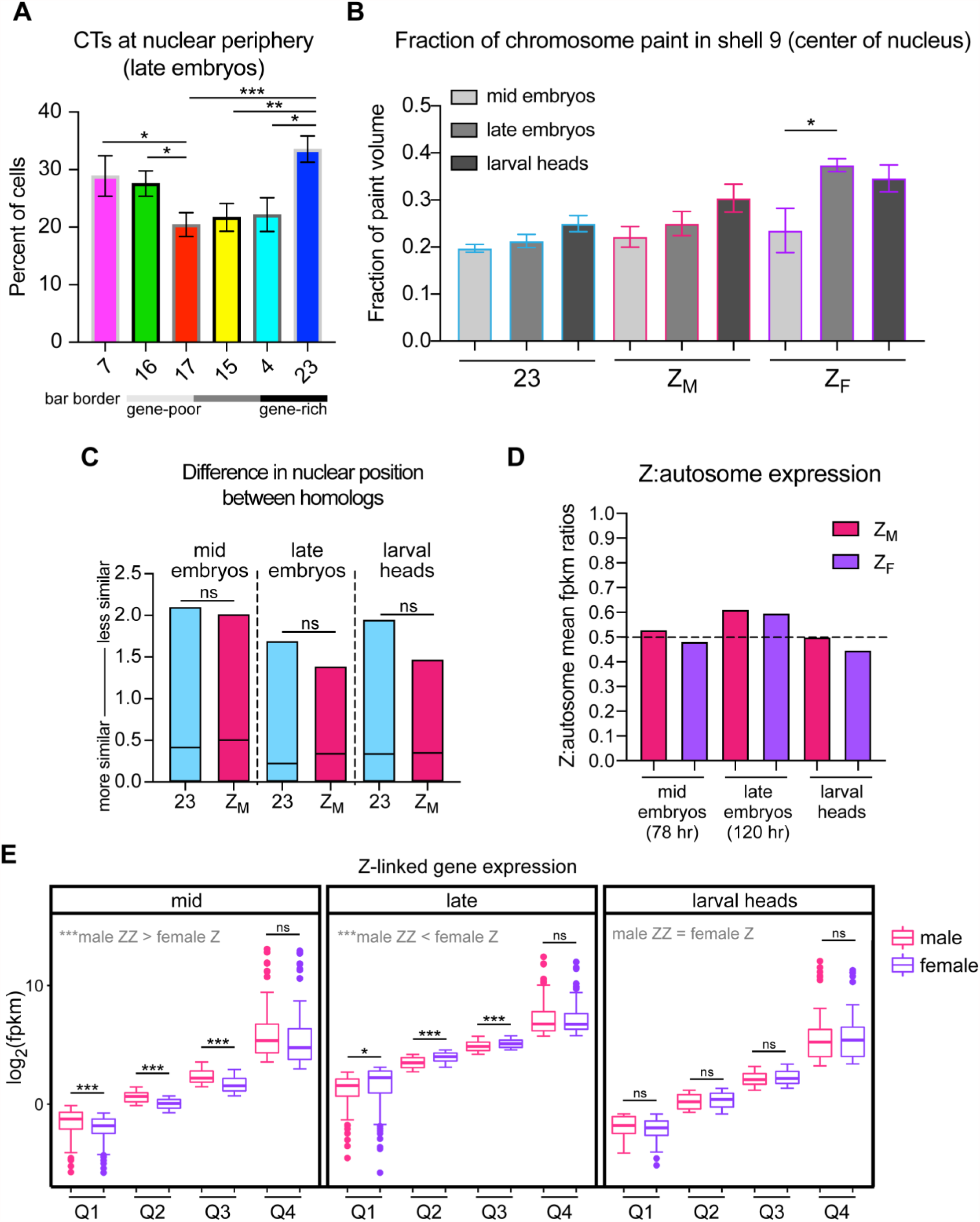
Female chZ is reposition toward the nuclear center and is upregulated in late embryos. A) Quantification of fraction of autosomes at nuclear periphery in late embryos (defined as having any volume in shells 1-3 of a shell analysis). Bars show average of biological replicates. Error bars show standard error of the mean. Statistics = unpaired t-tests. *p = 0.01 - 0.05, **p = 0.001 - 0.005, ***p < 0.001. Gene density of chromosomes is indicated by bar outlines, with darker outlines indicating higher gene density. B) Quantification of the fraction of CTs in the nuclear center (defined as having any volume in shell 9 of a shell analysis). Bars show the average between biological replicates. Error bars show standard error of the mean. *p=0.05; unpaired t-test. C) Quantification of interhomolog differences in nuclear position as measured by the distance from the nuclear edge (µm) for ch23 (blue) or chZ (pink) in male embryos or larvae. Mid-line = mean. Statistics = unpaired t-tests. D) Z:autosome mean fpkm expression ratios from RNA-seq data. A Z:autosome ratio of 0.5 indicates half as much expression on chZ compared to the autosomal average. E) Box and whisker plot showing the comparison between male and female quartile expression for Z-linked genes from h78 (mid/late), h120 (late), and head (larval head) data from Gopinath et al., 2019, where Q1 is the first quartile (lowest expression) and Q4 is the highest quartile (highest expression). Boxes indicate the interquartile range, and lines show the median. All unexpressed genes (fpkm = 0) were omitted. Statistics = Welch Two Sample t-test (two-sided). Late/mid: Q1 p=5.7e-05, Q2 p<2.22e-16, Q3 p=1.7e-15, Q4 p=0.11. Late: Q1 p=0.03, Q2 p=4.9e-15, Q3 p=5e-05, Q4 p=0.95. Larval head: Q1 p=0.29, Q2 p=0.26, Q3 p=0.21, Q4 p=0.72.

## References

1. T. Cremer, M. Cremer, Chromosome Territories. Cold Spring Harb Perspect Biol 2 (2010).

2. H. B. Sun, J. Shen, H. Yokota, Size-dependent positioning of human chromosomes in interphase nuclei. Biophys J 79, 184–190 (2000).

3. A. J. Fritz, N. Sehgal, A. Pliss, J. Xu, R. Berezney, Chromosome territories and the global regulation of the genome. Genes, Chromosomes and Cancer 58, 407–426 (2019).

4. L. Parada, T. Misteli, Chromosome positioning in the interphase nucleus. Trends Cell Biol. 12, 425–432 (2002).

5. L. F. Rosin, S. C. Nguyen, E. F. Joyce, Condensin II drives large-scale folding and spatial partitioning of interphase chromosomes in Drosophila nuclei. PLoS Genet. 14, e1007393 (2018).

6. L. Guelen, et al., Domain organization of human chromosomes revealed by mapping of nuclear lamina interactions. Nature 453, 948–951 (2008).

7. H. Pickersgill, et al., Characterization of the Drosophila melanogaster genome at the nuclear lamina. Nat. Genet. 38, 1005–1014 (2006).

8. B. van Steensel, A. S. Belmont, Lamina-Associated Domains: Links with Chromosome Architecture, Heterochromatin, and Gene Repression. Cell 169, 780–791 (2017).

9. B. van Steensel, S. Henikoff, Identification of in vivo DNA targets of chromatin proteins using tethered Dam methyltransferase. Nat Biotechnol 18, 424–428 (2000).

10. B. D. Towbin, et al., Step-wise methylation of histone H3K9 positions heterochromatin at the nuclear periphery. Cell 150, 934–947 (2012).

11. N. Briand, P. Collas, Lamina-associated domains: peripheral matters and internal affairs. Genome Biology 21, 85 (2020).

12. C. Vigouroux, et al., Nuclear envelope disorganization in fibroblasts from lipodystrophic patients with heterozygous R482Q/W mutations in the lamin A/C gene. J Cell Sci 114, 4459–4468 (2001).

13. H. J. Worman, Nuclear lamins and laminopathies. The Journal of Pathology 226, 316–325 (2012).

14. C. Ferrai, I. J. de Castro, L. Lavitas, M. Chotalia, A. Pombo, Gene Positioning. Cold Spring Harb Perspect Biol 2, a000588 (2010).

15. E. M. Blackwood, J. T. Kadonaga, Going the distance: a current view of enhancer action. Science 281, 60–63 (1998).

16. I. Chepelev, G. Wei, D. Wangsa, Q. Tang, K. Zhao, Characterization of genome-wide enhancer-promoter interactions reveals co-expression of interacting genes and modes of higher order chromatin organization. Cell Res. 22, 490–503 (2012).

17. F. Ciabrelli, G. Cavalli, Chromatin-Driven Behavior of Topologically Associating Domains. Journal of Molecular Biology 427, 608–625 (2015).

18. K. E. Cullen, M. P. Kladde, M. A. Seyfred, Interaction between transcription regulatory regions of prolactin chromatin. Science 261, 203–206 (1993).

19. D. A. Kleinjan, V. van Heyningen, Long-Range Control of Gene Expression: Emerging Mechanisms and Disruption in Disease. Am J Hum Genet 76, 8–32 (2005).

20. L. Gu, J. R. Walters, D. C. Knipple, Conserved Patterns of Sex Chromosome Dosage Compensation in the Lepidoptera (WZ/ZZ): Insights from a Moth Neo-Z Chromosome. Genome Biol Evol 9, 802–816 (2017).

21. I. Marín, M. L. Siegal, B. S. Baker, The evolution of dosage-compensation mechanisms. BioEssays 22, 1106–1114 (2000).

22. W. Jordan, L. E. Rieder, E. Larschan, Diverse Genome Topologies Characterize Dosage Compensation across Species. Trends Genet 35, 308–315 (2019).

23. S. Ercan, Mechanisms of X Chromosome Dosage Compensation. J Genomics 3, 1–19 (2015).

24. C. Grimaud, P. B. Becker, The dosage compensation complex shapes the conformation of the X chromosome in Drosophila. Genes Dev. 23, 2490–2495 (2009).

25. A. C. Lau, K. Nabeshima, G. Csankovszki, The C. elegans dosage compensation complex mediates interphase X chromosome compaction. Epigenetics & Chromatin 7, 31 (2014).

26. E. Splinter, et al., The inactive X chromosome adopts a unique three-dimensional conformation that is dependent on Xist RNA. Genes Dev 25, 1371–1383 (2011).

27. J. C. Chow, E. Heard, Nuclear Organization and Dosage Compensation. Cold Spring Harb Perspect Biol 2, a000604 (2010).

28. R. Sharma, et al., Differential spatial and structural organization of the X chromosome underlies dosage compensation in C. elegans. Genes Dev 28, 2591–2596 (2014).

29. L. Giorgetti, et al., Structural organization of the inactive X chromosome in the mouse. Nature 535, 575–579 (2016).

30. S. Ohno, Sex Chromosomes and Sex-Linked Genes (Springer Science & Business Media, 1967).

31. S. E. Albritton, et al., Sex-Biased Gene Expression and Evolution of the X Chromosome in Nematodes. Genetics 197, 865–883 (2014).

32. X. Deng, et al., Evidence for compensatory upregulation of expressed X-linked genes in mammals, Caenorhabditis elegans and Drosophila melanogaster. Nat. Genet. 43, 1179–1185 (2011).

33. X. Deng, J. B. Berletch, D. K. Nguyen, C. M. Disteche, X chromosome regulation: diverse patterns in development, tissues and disease. Nature Reviews Genetics 15, 367–378 (2014).

34. R. A. Veitia, F. Veyrunes, S. Bottani, J. A. Birchler, X chromosome inactivation and active X upregulation in therian mammals: facts, questions, and hypotheses. J Mol Cell Biol 7, 2–11 (2015).

35. B. S. Wheeler, et al., Chromosome-wide mechanisms to decouple gene expression from gene dose during sex-chromosome evolution. eLife 5, e17365 (2016).

36. H. Ellegren, et al., Faced with inequality: chicken do not have a general dosage compensation of sex-linked genes. BMC Biology 5, 40 (2007).

37. Y. Itoh, et al., Dosage compensation is less effective in birds than in mammals. J Biol 6, 2 (2007).

38. Y. Itoh, et al., Sex bias and dosage compensation in the zebra finch versus chicken genomes: General and specialized patterns among birds. Genome Res 20, 512–518 (2010).

39. B. Vicoso, D. Bachtrog, Lack of Global Dosage Compensation in Schistosoma mansoni, a Female-Heterogametic Parasite. Genome Biol Evol 3, 230–235 (2011).

40. J. B. Wolf, J. Bryk, General lack of global dosage compensation in ZZ/ZW systems? Broadening the perspective with RNA-seq. BMC Genomics 12, 91 (2011).

41. P. Julien, et al., Mechanisms and Evolutionary Patterns of Mammalian and Avian Dosage Compensation. PLOS Biology 10, e1001328 (2012).

42. S. Uebbing, A. Künstner, H. Mäkinen, H. Ellegren, Transcriptome Sequencing Reveals the Character of Incomplete Dosage Compensation across Multiple Tissues in Flycatchers. Genome Biol Evol 5, 1555–1566 (2013).

43. B. Vicoso, J. J. Emerson, Y. Zektser, S. Mahajan, D. Bachtrog, Comparative Sex Chromosome Genomics in Snakes: Differentiation, Evolutionary Strata, and Lack of Global Dosage Compensation. PLOS Biology 11, e1001643 (2013).

44. S. Chen, et al., Whole-genome sequence of a flatfish provides insights into ZW sex chromosome evolution and adaptation to a benthic lifestyle. Nature Genetics 46, 253–260 (2014).

45. G. Gopinath, K. Srikeerthana, A. Tomar, S. M. Ch. Sekhar, K. P. Arunkumar, RNA sequencing reveals a complete but an unconventional type of dosage compensation in the domestic silkworm Bombyx mori. Royal Society Open Science 4, 170261 (2017).

46. A. K. Huylmans, A. Macon, B. Vicoso, Global Dosage Compensation Is Ubiquitous in Lepidoptera, but Counteracted by the Masculinization of the Z Chromosome. Mol Biol Evol 34, 2637–2649 (2017).

47. T. Kiuchi, et al., A single female-specific piRNA is the primary determiner of sex in the silkworm. Nature 509, 633–636 (2014).

48. G. Smith, Y.-R. Chen, G. W. Blissard, A. D. Briscoe, Complete dosage compensation and sexbiased gene expression in the moth Manduca sexta. Genome Biol Evol 6, 526–537 (2014).

49. J. R. Walters, T. J. Hardcastle, Getting a full dose? Reconsidering sex chromosome dosage compensation in the silkworm, Bombyx mori. Genome Biol Evol 3, 491–504 (2011).

50. J. R. Walters, T. J. Hardcastle, C. D. Jiggins, Sex Chromosome Dosage Compensation in Heliconius Butterflies: Global yet Still Incomplete? Genome Biol Evol 7, 2545–2559 (2015).

51. S. Ercan, J. D. Lieb, C. elegans dosage compensation: a window into mechanisms of domain-scale gene regulation. Chromosome Res 17, 215–227 (2009).

52. V. Gupta, et al., Global analysis of X-chromosome dosage compensation. J Biol 5, 3 (2006).

53. K. Mita, et al., The genome sequence of silkworm, Bombyx mori. DNA Res. 11, 27–35 (2004).

54. Q. Xia, et al., A draft sequence for the genome of the domesticated silkworm (Bombyx mori). Science 306, 1937–1940 (2004).

55. T. D. King, et al., Recurrent losses and rapid evolution of the condensin II complex in insects. Mol. Biol. Evol. (2019) https://doi.org/10.1093/molbev/msz140.

56. M. Kawamoto, et al., High-quality genome assembly of the silkworm, Bombyx mori. Insect Biochemistry and Molecular Biology 107, 53–62 (2019).

57. L. F. Rosin, J. Gil, I. A. Drinnenberg, E. P. Lei, Oligopaint DNA FISH as a tool for investigating meiotic chromosome dynamics in the silkworm, Bombyx mori. bioRxiv, 2021.04.16.440181 (2021).

58. E. H. Finn, et al., Extensive Heterogeneity and Intrinsic Variation in Spatial Genome Organization. Cell 176, 1502–1515.e10 (2019).

59. L. Gu, et al., Dichotomy of Dosage Compensation along the Neo Z Chromosome of the Monarch Butterfly. Curr Biol 29, 4071–4077.e3 (2019).

60. A. P. Senaratne, et al., Formation of the CenH3-Deficient Holocentromere in Lepidoptera Avoids Active Chromatin. Current Biology (2020) https://doi.org/10.1016/j.cub.2020.09.078 (December 9, 2020).

61. T. Ando, T. Kojima, H. Fujiwara, Dramatic changes in patterning gene expression during metamorphosis are associated with the formation of a feather-like antenna by the silk moth, Bombyx mori. Developmental Biology 357, 53–63 (2011).

62. R. Sharma, P. Meister, Linking dosage compensation and X chromosome nuclear organization in C. elegans. Nucleus 6, 266–272 (2015).

63. P. T. Chuang, D. G. Albertson, B. J. Meyer, DPY-27:a chromosome condensation protein homolog that regulates C. elegans dosage compensation through association with the X chromosome. Cell 79, 459–474 (1994).

64. I. Bag, et al., M1BP cooperates with CP190 to activate transcription at TAD borders and promote chromatin insulator activity. bioRxiv, 2020.10.27.357533 (2021).

65. D. Chen, R. K. Dale, E. P. Lei, Shep regulates Drosophila neuronal remodeling by controlling transcription of its chromatin targets. Development 145 (2018).

66. B. J. Beliveau, et al., OligoMiner provides a rapid, flexible environment for the design of genome-scale oligonucleotide in situ hybridization probes. Proc. Natl. Acad. Sci. U.S.A. 115, E2183–E2192 (2018).

67. B. D. Fields, S. C. Nguyen, G. Nir, S. Kennedy, A multiplexed DNA FISH strategy for assessing genome architecture in Caenorhabditis elegans. eLife 8 (2019).

68. S. C. Nguyen, E. F. Joyce, “Programmable Chromosome Painting with Oligopaints” in Imaging Gene Expression: Methods and Protocols, Methods in Molecular Biology., Y. Shav-Tal, Ed. (Springer New York, 2019), pp. 167–180.

69. J. Ollion, J. Cochennec, F. Loll, C. Escudé, T. Boudier, TANGO: a generic tool for high-throughput 3D image analysis for studying nuclear organization. Bioinformatics 29, 1840–1841 (2013).

70. B. Bushnell, J. Rood, E. Singer, BBMerge - Accurate paired shotgun read merging via overlap. PLoS One 12, e0185056 (2017).

71. A. M. Bolger, M. Lohse, B. Usadel, Trimmomatic: a flexible trimmer for Illumina sequence data. Bioinformatics 30, 2114–2120 (2014).

72. B. Langmead, S. Salzberg, Fast gapped-read alignment with Bowtie 2 | Nature Methods. Nature Methods (2012) https://doi.org/10.1038/nmeth.1923 (April 19, 2021).

73. H. Li, et al., The Sequence Alignment/Map format and SAMtools. Bioinformatics 25, 2078–2079 (2009).

74. Y. Liao, G. K. Smyth, W. Shi, featureCounts: an efficient general purpose program for assigning sequence reads to genomic features. Bioinformatics 30, 923–930 (2014).

75. Y. Suetsugu, et al., Large Scale Full-Length cDNA Sequencing Reveals a Unique Genomic Landscape in a Lepidopteran Model Insect, Bombyx mori. G3 (Bethesda) 3, 1481–1492 (2013).

